# Maximized redundant and synergistic information transfers predict the rise in the output gene expression noise in a generic class of coherent type-1 feed-forward loop networks

**DOI:** 10.1101/2021.09.12.459930

**Authors:** Md Sorique Aziz Momin, Ayan Biswas

## Abstract

We apply the partial information decomposition principle to a generic coherent type-1 feed-forward loop (C1-FFL) motif with tunable direct and indirect transcriptional regulations of the output gene product and quantify the redundant, synergistic, and unique information transfers from the regulators to their target output species. Our results which are obtained within the small-noise regime of a Gaussian framework reveal that the redundant and synergistic information transfers are antagonistically related to the output noise. Most importantly, these two information flavors are maximized prior to the minimization and subsequent growth of the output noise. Therefore, we hypothesize that the dynamic information redundancy and synergy maxima may possibly be utilized as efficient statistical predictors to forecast the increasing trend of the fluctuations associated with the output gene expression dynamics in the C1-FFL class of network motifs. Our core analytical finding is supported by exact stochastic simulation data and furthermore validated for a diversified repertoire of biologically plausible parameters. Since, the output gene product serves essential physiological purposes in the cell, a predictive estimate of its noise level is supposed to be of considerable biophysical utility.

## I. INTRODUCTION

Cells maintain an intricate relationship with their environments where they come across different stimuli such as usable nutrients, detrimental molecules, external stresses (e.g., heat-shock and osmotic-pressure), etc. In order to survive in these convoluted circumstances, cells respond to the incoming signals by producing appropriate proteins/enzymes at the right time and in economic amounts to achieve desired physiological goals with appreciable precision. A typical *E. coli* and yeast cells contain roughly 4 × 10^6^ and 4 × 10^9^ proteins, respectively, with individual types of protein numbered in 𝒪 (100) copies [1–3]. The instructions for synthesizing this large number of different proteins come from different genes: the coding regions of DNA. The information processing in between environmental signals and genes is mediated by a special type of catalytic protein, namely, transcription factor (TF). At first, the inducer signal activates certain TFs which, upon binding to the regulatory promoter regions of their enslaved genes, increase (activation) or decrease (repression) the transcription rates. This is done by altering the binding probabilities of the target genes with the RNA polymerase. Thus, the production rates of mRNAs and consequently that of the end-products (proteins/enzymes) are modified at par with the cellular necessities. Some of these gene products can act as TFs thereby regulating other genes both up and downstream along the signaling pathway. This TF-gene interaction which forms an arrayed structure is generally known as the gene transcription regulatory network (GTRN) [2].

The GTRN of an organism, in terms of its size and activities, is very complex in nature and contains different architectural patterns, e.g., cascades, feed-forward and feed-back loops, etc. Some of these network patterns, often of small sizes, are over-represented in the GTRN of a particular organism than in an equivalent randomized network. These recurring small networks are known as the network motifs [4]. Associating three genes/gene products as different nodes and their activatory interactions as the inter-node edges, there appears an ensemble of thirteen three-node random subgraphs. Out of these different possible network configurations, the C1-FFL is found to appear significantly in the GTRNs of *E. coli*, yeast, and many other higher organisms. Hence, for these GTRNs, the C1-FFL is qualified as the only network motif. Within this motif, TF “S” activates another TF “X” whereas both of them activate the production from the output gene “Y”. Here, for simplicity, we use the same notations for the biochemical species (genes and gene products) and their population levels. Their meanings should be inferred depending upon the context of their usage. Therefore, the production of gene product Y is accomplished through two parallel regulatory paths: direct regulation (S → Y) and another indirect regulation (S → X → Y). In the terminology of networks, the former regulatory process forms a one-step cascade whereas the latter constitutes a two-step cascade. In the absence of an edge between X and Y, one obtains another additional one-step process: S → X apart from the previously mentioned S → Y. The resulting pattern out of these two one-step branches is: Y ← S → X otherwise known as the fan-out network. For a generic C1-FFL with tunable direct and indirect regulatory controls over its output gene, the two-step cascade and fan-out structure may be considered as two of its extreme realizations where the direct and indirect decoding rate, respectively, are zero! Therefore, except for these two special patterns, a typical C1-FFL motif is capable of decoding the upstream signal through two parallel regulatory pathways. The decoding process is perfect when their individual shares in the output gene expression level are equal in amounts.

This paper intends to explore the information processing capabilities of a generic C1-FFL motif by keeping the interplay of its direct and indirect decoding strengths in perspective. To be specific, our objective is to build a multivariate information-theoretic framework to compare different types of the intergenetic information transfer and their relationships with the output gene expression noise for the generalized C1-FFL class of motifs. According to our network modification scheme which will be formally introduced in Sec. II C, each of these C1-FFL configurations are genotypically different by means of different direct and indirect production rates (decoding strengths) of the parallel regulatory pathways. In contrast, our biochemical construct envisages a situation in which all the inter-convertible structures of the C1-FFL type may manage to maintain fixed expression levels of the three constituent genes. This constraint on the gene products’ abundances implies that different C1-FFLs are equitable on phenotypic footing although being distinct in the underlying genotypic spaces. To add a biological perspective to our current information-theoretic question, it is imperative to mention that the C1-FFL motif performs essential physiological functions in living systems. To exemplify, C1-FFL employing a SUM signal integration logic is operational in making flagellar motor proteins in *E. coli* [5]. The network modification scheme in our work is somewhat broadly inspired, albeit on a theoretical front, by the foundational experiments performed on the *arabinose* utilization system in *E. coli*. There, Mangan and colleagues have compared a C1-FFL module with one of its extreme network instances, namely, the simple regulation which is otherwise known as the fan-in network. Within the fan-in network, the two regulators do not interact transcriptionally with each other [6] and interestingly, just like the two-step cascade and fan-out network, the fan-in pattern is also a member of the previously mentioned ensemble of thirteen three-node random networks. Although, the biological impact of our hypothesized construct is further subject to experimental scrutiny, the prescribed theoretical model is a rich ground to explore the information processing implications of plausible modular topologies within the broad C1-FFL family.

The biochemical kinetics associated with the synthesis and degradation of different gene products are afflicted with noise in primarily two ways: (i) due to the low copy numbers of the biomolecules participating in various chemical reactions and (ii) fluctuations of global origin flowing downstream to the enslaved genes. Therefore, noise is an inherent element to living systems, specifically in the context of a single cell, and thus becomes an indispensable feature of cellular functionalities at both local and global levels [7–11]. To coherently describe the noisy gene expression dynamics at the single-cell level, one needs to bring in the tools and techniques of stochastic processes. An information-theoretic framework [12–14] is one of such routes which coarse-grains the biochemical signaling process as a noisy input/output communication channel. The application of information theory is diversified as it spans the domain of neuroscience, gene transcription regulation as well as intercellular communication, to name a few [15–26]. In this paper, we study the effects of interspecies interactions within an ensemble of C1-FFLs with tunable direct and indirect decoding strengths on the efficient maintenance of the directional information flow. Transfer entropy (TE) offers us the mathematical machinery to quantify this information dynamics which detect the asymmetric statistical dependencies between different biochemical species in C1-FFL patterns [27, 28]. In recent years, applications of the TE-based measures have successfully generated new insights in a number of physical and physiological systems, e.g., Ising models, epileptic brains, *HeLa* cell-cultures, etc. [29–31]. On the other side, the scope of measuring entropy transfer in the GTRN motifs is still unexplored to a significant extent. The current paper is intended to make progress in this uncharted domain of research. Such an information-theoretic construct fundamentally involves measuring correlations between gene expression levels from two nearby time points at steady state in contrast with single-time-point measurements. The metric of TE goes beyond measuring the nonlinear correlation between different gene products and quantifies the extent to which the information from the past states of a set of source species can predict the future state of the target species. Thus, TE provides us with a mechanistic understanding of the control over the fluctuations in the target gene expression using the information sources. For our model C1-FFL, as will be demonstrated in Sec. III A, the information transferred from the regulators to their enslaved gene product often comes from unique sources. Other types of information flow may involve either the master- or the co-regulator depending upon the relative values of their information transfer capacities. Additionally, the regulators may even complement each other to transfer information. Here, we are concerned with the question: How different types of information transfer are interrelated with the output noise? The dynamical C1-FFL model will be introduced in Sec. II A. To establish the required analytical framework in discrete-time domain, we apply a stationary multivariate vector autoregressive (MVAR) process [31–33] as deliberated in the Appendix which also contains the details of the stochastic simulation method. The information-theoretic measures will be formally introduced in Sec. II B. The scheme for the architectural modification and the rationale behind our choices of parameters are contained in Sec. II C. Sec. III A provides our main analytical results supported by exact stochastic simulation data. Secs. (III B-III E) provide further analytical support for our core findings within a broad regime of parametric realization. Finally, we conclude our article with a summary of our findings in Sec. IV.

## II. THE MODEL

### A. Gene expression dynamics in a coarse-grained C1-FFL motif

Using the Langevin formalism [24, 34, 35], we model the dynamics of the populations (random variables) of the three gene products (S, X, and Y) which constitute the C1-FFL motif. These population (gene expression) levels are expressed in the unit of copy numbers in an effective cellular volume of magnitude unity. The set of dynamical equations is as follows:

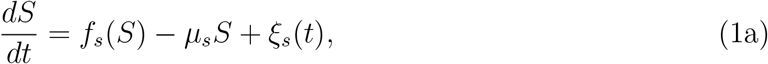

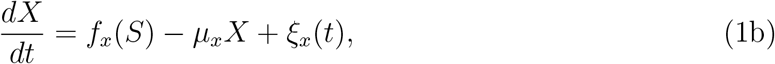

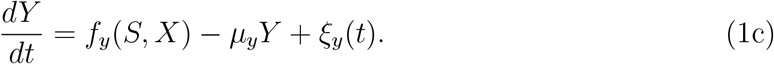

*f*_*s*_, *f*_*x*_(*S*), and *f*_*y*_(*S, X*) are the production/synthesis rates of S, X, and Y, respectively. These input rate functions may depend upon the regulators’ populations in a (non)linear fashion. Usage of the Hill function to model transcriptional regulation is a standard practice supported by experimental data [2, 16, 36–38]. To exemplify in the present model, the synthesis rate for S → X is taken: *f*_*x*_(*S*) = *k*_*sx*_[*S/*(*K*_*sx*_ + *S*)] where *k*_*sx*_ is the synthesis rate parameter interpreted as the maximal production rate from the gene promoter of X. The remaining part in *f*_*x*_ describes the promoter occupancy probability of gene X by the activated TF S. The activation coefficient *K*_*sx*_ is numerically equivalent to the copy number of the activated TF S which is needed to reach the half-maximal expression of X [2]. The production rate for the reaction channel: *ϕ* → S is taken to be independent of any TF population external to the three-node subgraph and hence, we have *f*_*s*_ = *k*_*s*_. The combined output production rate considering an additive logic operating upon the direct and indirect regulatory pathways is *k*_*sy*_[*S/*(*K*_*sy*_ + *S*)] + *k*_*xy*_[*X/*(*K*_*xy*_ + *X*)]. *k*_*sy*_ (*k*_*xy*_) is the production rate parameter for S → Y (X → Y) whereas *K*_*sy*_ (*K*_*xy*_) is the corresponding activation coefficient. The degradation rates of S, X, and Y are modelled as linearly proportional to their corresponding populations, i.e., *µ*_*s*_*S, µ*_*x*_*X*, and *µ*_*y*_*Y*, respectively, where *µ*_*s*_, *µ*_*x*_, and *µ*_*y*_ are the related degradation rate parameters. To represent the inherently stochastic synthesis and degradation reactions, the Langevin description includes the Gaussian-distributed white noise terms *ξ*_*s*_(*t*), *ξ*_*x*_(*t*), and *ξ*_*y*_(*t*) for species S, X, and Y, respectively. Their statistical properties are as follows [39–42]: ⟨*ξ*_*z*_(*t*)⟩= 0, ⟨*ξ*_*z*_(*t*)*ξ*_z′_(*t′*)⟩ = ⟨|*ξ*_*z*_(*t*)|^2^⟩*δ*_*zz*′_*δ*(*t* − *t′*) with {*z, z′*} ∈ {*s, x, y*} being the respective indices for gene products S, X, and Y. *δ*_*zz*′_and *δ*(*t* − *t′*) represent the Kronecker- and Dirac-delta functions, respectively. ⟨…⟩ indicates steady-state ensemble-averages whereas *t* and *t′* stand for two different time points. The steady-state ensemble-averaged noise strength is defined as ⟨|*ξ*_*z*_(*t*)|^2^⟩= ⟨*f*_*z*_(*Z*)⟩+ *µ*_*z*_ ⟨*Z*⟩= 2*µ*_*z*_ ⟨*Z*⟩where, *Z* ∈ {*S, X, Y*} are the populations of different gene products. The last equality comes from the steady-state ensemble-averaged versions of Eqs. (1a-1c). The steady-state expressions of the population levels of the gene products are: ⟨*S*⟩ = *k*_*s*_*/µ*_*s*_, ⟨*X*⟩ = (*k*_*sx*_*/µ*_*x*_) ⟨*f*_*x*_(*S*)⟩ ≈ (*k*_*sx*_*/µ*_*x*_)*f*_*x*_(⟨*S*⟩) = (*k*_*sx*_*/µ*_*x*_) [⟨*S*⟩*/*(*K*_*sx*_ + ⟨*S*⟩)]. The approximation is reasonable within the small-noise regime [24]. Similarly, ⟨*Y*⟩ = ⟨*Y* (*S*)⟩+ ⟨*Y* (*X*)⟩ where ⟨*Y* (*S*)⟩ ≈ (*k*_*sy*_*/µ*_*y*_) [⟨*S*⟩*/*(*K*_*sy*_+ ⟨*S*⟩)] and ⟨*Y* (*X*)⟩ ≈ (*k*_*xy*_*/µ*_*y*_) [⟨*X*⟩*/*(*K*_*xy*_+ ⟨*X*⟩)]. Evidently, ⟨*Y* (*S*)⟩ and ⟨*Y* (*X*)⟩ are the steady-state ensemble-averaged populations of the output gene product which are synthesized by the master- and co-regulator, respectively. The units for the degradation rate parameters are min^−1^. The synthesis rate parameters are expressed in the units of c.n.×min^−1^ where c.n. stands for copy number (per an effective cellular volume of magnitude unity). The activation coefficients have the units of c.n. as in the case for population levels. Coming back to the random forcing terms (*ξ*_*z*_(*t*)), we have used Gaussian noise with zero means and zero cross-species correlations as well as zero temporal correlations. These are reasonable assumptions for coarse-grained transcription networks which incorporate birth-death types of reactions like our present model [24]. At steady state, noise strengths are conceived to originate equally from the stochastic synthesis and degradation events of each of the biochemical species. Besides, usage of constant steady-state noise strengths is an approximation adopted to ease the analytical tractability of the forthcoming computations of the (co)variances of various gene products [see the Appendix]. There, we will also discuss the effect of the time-discretization of Eqs. (1a-1c) on the noise strengths.

### B. Information-theoretic measures

While explicitly dealing with the temporal structure of gene expression data, the one-time-point/static mutual information (MI) can be upgraded to evaluate the association between two random variables from two different time points, e.g., *Y*_*t*_ and *Y*_*t*+1_. This enables us to quantify the dynamics of MI. In both the static and dynamic cases, MI may involve more than two random variables corresponding to distinct biochemical species. By judiciously combining two- and three-variable two-time-point MIs together, we can alleviate the non-directional aspect of the static information. The resulting metric of TE quantifies the information transfer from the input to the output level of an information channel. The joint effect of two source species in transferring information to a single target species is also possible to compute using the multivariate TE. In our present case, TE may be used to predict the future state of the output (i.e., *Y*_*t*+1_) using the combined knowledge of its present state (*Y*_*t*_) and that of another source species (e.g., *S*_*t*_) beyond what can be predicted by *Y*_*t*_ alone [27, 28, 43, 44]. Therefore, TE from S to Y is expressed as:

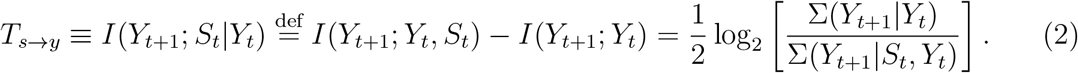

Here, *I*(⋯ ; ⋯) are various dynamic MIs whereas *I*(⋯ | ⋯) represent different conditional two-time-point MIs. The partial variances Σ(⋯ | ⋯) are computed using Eqs. (A.10a, A.10b) given in the Appendix. The last equality in terms of the time-lagged partial variances is naturally established for the present Gaussian system [31, 32, 45]. Similarly, TE from X to Y is computed using:

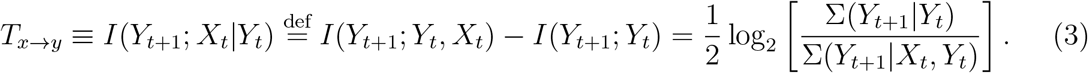

Lastly, TE due to the joint action of S and X targeting Y is the following:

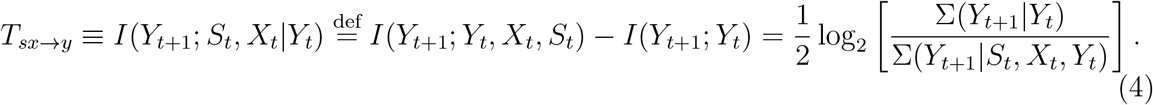

To reiterate, the TE metrics: *T*_*s*→*y*_, *T*_*x*→*y*_, and *T*_*sx*→*y*_ measure the information contents transferred from S to Y, from X to Y, and jointly from S and X to Y, respectively.

According to the prescription of *Partial information decomposition* (PID), *T*_*sx*→*y*_ is dissected into four distinct/independent non-negative and non-overlapping information transfer elements. Unique information *U*_*s*→*y*_ is transferred to Y from S alone. Similarly, unique information *U*_*x*→*y*_ is directed to Y only from X. Redundant information transfer *R*_*sx*→*y*_ is due to that regulator which transmits a lesser amount of information in comparison with the other information source. On the other hand, *S*_*sx*→*y*_ is the synergistic information transmitted complementarily by the pair of sources. As the name suggests, synergistic information is the amount of information processed by the source variables/species operating in unison while predicting their target. The set of PID elements are related to the TE metrics by the following set of equations [31, 32, 46–48]:

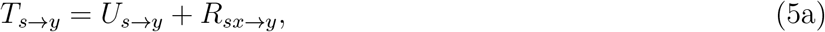

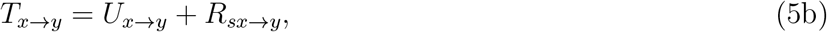

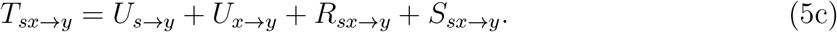

In the transfer entropy space, *T*_*s*→*y*_ and *T*_*x*→*y*_ are embedded within *T*_*sx*→*y*_. The non-overlapping portions of *T*_*s*→*y*_ and *T*_*x*→*y*_ contribute the unique information transfers, i.e., *U*_*s*→*y*_ and *U*_*x*→*y*_ whereas the common (overlapping) space provides *R*_*sx*→*y*_. *S*_*sx*→*y*_ lies beyond the combined space of *T*_*s*→*y*_ and *T*_*x*→*y*_ and is contained within the space of *T*_*sx*→*y*_ [31]. To completely specify the PID elements in Eqs. (5a-5c), the redundant information transfer is defined as the minimum of the single-source-species TEs considering a unique target, i.e., *T*_*s*→*y*_ and *T*_*x*→*y*_. Thus, the formula to evaluate the redundant information transfer stands as [32]:

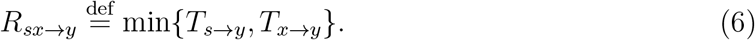

Finally, using the formalism of the *minimum mutual information* PID (MMI PID), i.e., Eqs. (5a-5c, 6), we can quantify the unique, synergistic, and redundant information transfers at steady state for the C1-FFL class of motifs. These working formulae are common for our (approximate) analytical and (exact) stochastic simulation-based data. Due to the usage of base “2” for the logarithm functions, the TEs and the PID elements are measured in the units of “bits”. Besides, we compute the noise level in the output gene expression using the concept of *coefficient of variation* (CV). The metric of interest is 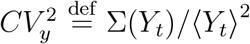. We use Eq. A.7f to analytically evaluate the output noise which is also measured exactly from the stochastic simulation. The output noise has the units of c.n.^−1^. Sec. III A presents these datasets which form the basis of our conjecture regarding the connection between the i/o information transfers and the output gene expression noise in the C1-FFL modular configurations.

### C. Architectural modification scheme for the C1-FFL class of networks

In one of the extreme configurations of the generic C1-FFL where the direct transcriptional regulation is absent (*k*_*sy*_ = 0 ⇒ ⟨*Y* (*S*)⟩= 0), Y is entirely produced by the TF X (i.e., ⟨*Y*⟩ = ⟨*Y* (*X*)⟩). Now, increasing ⟨*Y* (*S*)⟩ by one copy at a time by means of an appropriate increment in *k*_*sy*_ and simultaneously decreasing ⟨*Y* (*X*)⟩ by the same amount through lowering *k*_*xy*_, we reach a set of production rates where all of Y is produced by S alone (i.e., ⟨*Y*⟩ = ⟨*Y* (*S*)⟩ with *k*_*xy*_ and ⟨*Y* (*X*)⟩ = 0). In this other extreme realization of the C1-FFL motif, TF X does not share an edge with the output species Y. Hence, in between these two extreme points in the C1-FFL configuration space, schematized in Fig. 1, we have a range of intermediate structures with non-zero ⟨*Y* (*S*)⟩ and ⟨*Y* (*X*)⟩ (⟨*Y* (*S*)⟩ = 1 → 99 and ⟨*Y* (*X*)⟩ = 99 → 1 subject to the constraint: ⟨*Y*⟩ = ⟨*Y* (*S*)⟩ + ⟨*Y* (*X*)⟩= 100). Hence, the ratio of the direct production level to the total production level of the output gene product serves as an ideal normalized metric to indicate the inter-transmutations of different varieties of the C1-FFL network modules. In Sec. III, we will analyze the PID and noise elements with respect to this transformed tuning parameter: ⟨*Y* (*S*)⟩ ^*f*^ =: ⟨*Y* (*S*)⟩/⟨*Y*⟩. ⟨*Y* (*S*)⟩ = ⟨*Y* (*X*)⟩= ⟨*Y*⟩ */*2, or ⟨*Y* (*S*)⟩^*f*^ = 0.5 corresponds to a perfect C1-FFL structure with equally strong direct (S → Y) and indirect (X → Y) decoding paths.

**FIG. 1.**
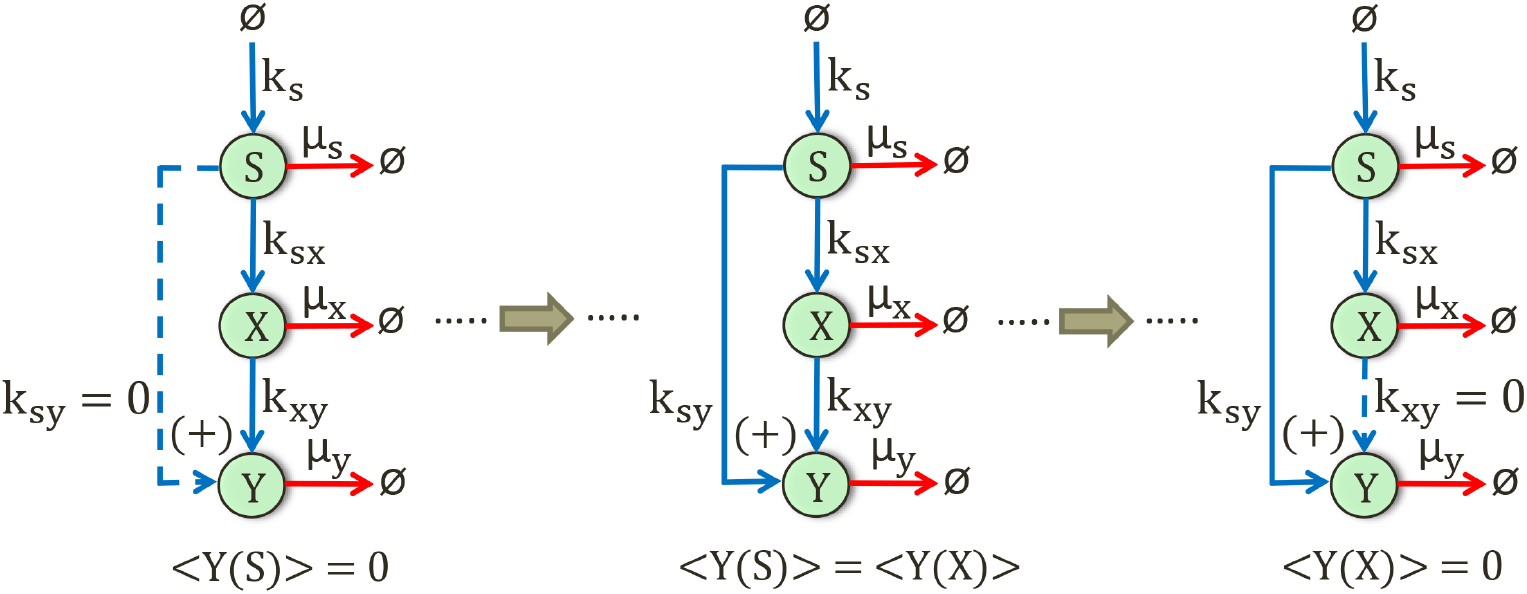
A schematic wiring diagram of a generic C1-FFL motif enabled with an additive logic (denoted by “+”) and tunable direct and indirect regulatory controls over the output gene product. The blue and red arrows designate synthesis (due to transcriptional activation) and degradation reactions of gene products, respectively. *k*_.._ and *µ*_.._ are the corresponding reaction rate parameters [see Sec. II(A)]. The extreme left and right configurations are special types of a generic C1-FFL motif where one of the regulatory branches leading up to the output species is absent. These non-existent interactions are marked by dashed blue lines with *k*_*sy*_ (⟨*Y* (*S*)⟩) or *k*_*xy*_ (⟨*Y* (*X*)⟩) = 0. The middle topology depicts a perfect C1-FFL with ⟨*Y* (*S*)⟩ = ⟨*Y* (*X*)⟩. *k*_*sy*_ (*k*_*xy*_) is increased (decreased) in a way so that ⟨*Y* (*S*)⟩ (⟨*Y* (*X*)⟩) increases (decreases) by one copy at a time thereby keeping the total population ⟨*Y*⟩ = ⟨*Y* (*S*)⟩ + ⟨*Y* (*X*)⟩ fixed. Hence, the entire network modification scheme reveals a variety of C1-FFLs each of which is characterized by unique sets of {*k*_*sy*_, *k*_*xy*_} or equivalently {⟨*Y* (*S*)⟩, ⟨*Y* (*X*)⟩}.

The strength of interaction between each of the target promoters of S and X on the output gene Y and the corresponding RNA polymerase is designated by the maximal production rates, i.e., *k*_*sy*_ and *k*_*xy*_, respectively. These synthesis rate parameters form part of the genotypic space of the generic C1-FFL whereas the output abundance level (⟨*Y*⟩) is its phenotypic characteristic. The latter is essentially interlinked with the benefits derived from this motif by a single cell. Therefore, a fixed value of ⟨*Y*⟩ signifies a many-to-one mapping from the genotypic to the phenotypic space. By maintaining fixed steady-state population levels: ⟨*S*⟩ = ⟨*X*⟩ = ⟨*Y*⟩ = 100, we ensure that different configurations of the generic C1-FFL type are compared on an equal footing [2]. The adoption of steady states to analyze the information transfer and noise management is motivated by the fact that the steady-state populations of the biochemical species are responsible for their optimal functioning in the single cell [2]. Fixed regulator abundances also imply that the TF-promoter interactions with the output gene (strengths of the Hill functions) are fixed. If we relax this constraint of population constancy and instead increase ⟨*S*⟩ and ⟨*X*⟩, the regulator-level intrinsic variability/noise ({⟨*S*⟩^−1^, ⟨*X*⟩^−1^}) which act like extrinsic noise for their enslaved gene products will drop. The reduction in flow of fluctuations will automatically result in reduced magnitudes of the entropy transfers and PID metrics. Since, the TE-based measures typically have low numerical values, we remain cautious about further lowering of their magnitudes. Keeping in mind the translocation of TF proteins from the cytoplasm to the nucleus in eukaryotic cells, the population levels of all the gene products are expressed in the units of molecules per effective cellular volume of magnitude unity. The degradation rate parameters considered for our main set of results are: *µ*_*s*_ = 0.1, *µ*_*x*_ = 0.5, and *µ*_*y*_ = 5.0 all in the units of min^−1^. With upstream regulators more stable than their downstream targets (*µ*_*s*_ *< µ*_*x*_ *< µ*_*y*_), we ensure sufficient flow of fluctuations and avoid the risk of insignificant PID values. In Sec. III C, we will examine the effects of a tunable input degradation rate parameter on our metrics of interest. The Hill coefficient *n* = 1 is used to avoid strong nonlinear effects. Activation coefficients are set at *K*_*sx*_ = *K*_*sy*_ = ⟨*S*⟩ and *K*_*xy*_ = ⟨*X*⟩ in units identical with that of the population levels. This type of choice denotes a particular operating point on the activation curves of X and Y where their promoter regions are half-filled by their respective regulators. Quantitatively speaking, the approximated steady-state values of the Hill functions thus become 0.5 and hence this type of regulatory state is known as the half-maximal activation [2]. If the activation coefficients are taken to be lower than the previously mentioned values, the input functions start to get steeper/nonlinear. The alternative choice, i.e., activation coefficients being higher than the population levels, make the input functions more linear but experimental wisdom suggests that the transcriptional control starts to mount with sufficient efficacy at the half-maximal point on the activation curve [2]. Considering these trade-offs, we present our main analytical and stochastic simulation results by taking into account the half-maximal activation. In Sec. III D, we will consider variations of activation coefficients around their half-maximal points. Since, the principle of PID is applicable for either linear or linearized systems, the values of *n* and *K*_*···*_ should be chosen in a way so that the input functions become sufficiently linearized. The activation coefficient is represented as the ratio of the off and on rate parameters of TF-promoter interaction, i.e., the ratio of rate parameters corresponding to the backward and forward reactions between a certain TF and its target promoter, forming the TF-promoter complex. For 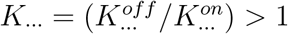, it is evident that 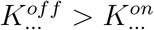. On the other side, the binding energy is quantified as *E*_*b*_ = −*k*_*B*_*T* ln *K*_*···*_ [3, 17]. Therefore, *K*_*···*_ *>* 1 suggests that *E*_*b*_ ≠ 0 which is an indirect signature of the underlying out-of-equilibrium dynamics of the system. It is to be pointed out that there exists a clear separation of timescales between different stages of gene transcription regulation. For instance, the inducer-dependent activations of the TFs and the formations of the TF-promoter complexes occur at a relatively faster timescales with respect to the time taken by the gene products to achieve their steady-state population levels. Taking this into consideration, our present coarse-grained dynamical model portrays a suitably approximated equilibrium picture of the actual nonequilibrium character of the biochemical kinetics [2]. The synthesis rate parameters *k*_*s*_, *k*_*sx*_, *k*_*sy*_, and *k*_*xy*_ are determined using the steady-state versions of Eqs. (1a-1c) along with the steady-state approximate forms of the Hill functions and the fixed steady-state population constraints. In Sec. III E, we will loosen this constraint of identical population levels of the regulators and the output gene product and examine the resulting effects on the PID measures and the output noise.

## III. RESULTS AND DISCUSSION

### A. Maxima of the synergistic and redundant information transfers indicate the impending growth of the output noise

In Fig. 2(a), we observe that with increasing direct production strength or ⟨*Y* (*S*)⟩ ^*f*^, both the redundant and synergistic information transfers from the regulators increase and maximize for a nearly balanced C1-FFL (⟨*Y* (*S*)⟩ ^*f*^ = 0.53). Since, the quantification of *R*_*sx*→*y*_ and *S*_*sx*→*y*_ involves both of the regulators, one can interpret their simultaneous maximization in terms of approximately equal amounts of ⟨*Y* (*S*)⟩ and ⟨*Y* (*X*)⟩. It is the direct and indirect productions of the output gene product through which the information processed at the interregulator level flows to the terminal species. We also notice that for this near-perfect configuration of the C1-FFL, *U*_*s*→*y*_ takes its first non-zero value whereas *U*_*x*→*y*_ turns zero. This further demonstrates that the two types of the unique information transfers can only exist independent of each other. For this C1-FFL topology, the regulators predominantly convey the redundant and synergistic information flavors to their predicted biochemical species. The decline in ⟨*Y* (*X*)⟩ level during the growth of redundancy and synergy explains the diminishing trend of information uniquely transferred by the co-regulator X. Similar logic gives out meaning to the trend of *U*_*s*→*y*_. The dominance of *R*_*sx*→*y*_ over *S*_*sx*→*y*_ as documented in Fig. 2(a) is also significantly observed in other small networks under the control of different tunable biochemical parameters [34]. Interestingly, the regulators continue to transfer the redundant information even when one of the regulatory pathways that control the output gene expression is absent but this is not the case with the synergistic information transfer. The numerical values of the PID elements have specific physical meanings attached to them and may be understood by taking into account the physical interpretation attributed to the concept of information, in general [49, 50]. Regulator S (X) transferring *U*_*s*→*y*_ (*U*_*x*→*y*_) bits of information to Y signifies that the predictor state *S*_*t*_ (*X*_*t*_) uniquely reduces the uncertainty of the output gene expression state *Y*_*t*+1_ by an extra factor of 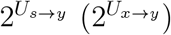, on average, with respect to the datum line contribution from *Y*_*t*_ to *Y*_*t*+1_. Similarly, the predictor states (*S*_*t*_ and *X*_*t*_) reduce the entropy of *Y*_*t*+1_ on a shared (complementary) basis by an extra factor of 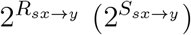, with respect to the reference entropy reduction done by *Y*_*t*_.

**FIG. 2.**
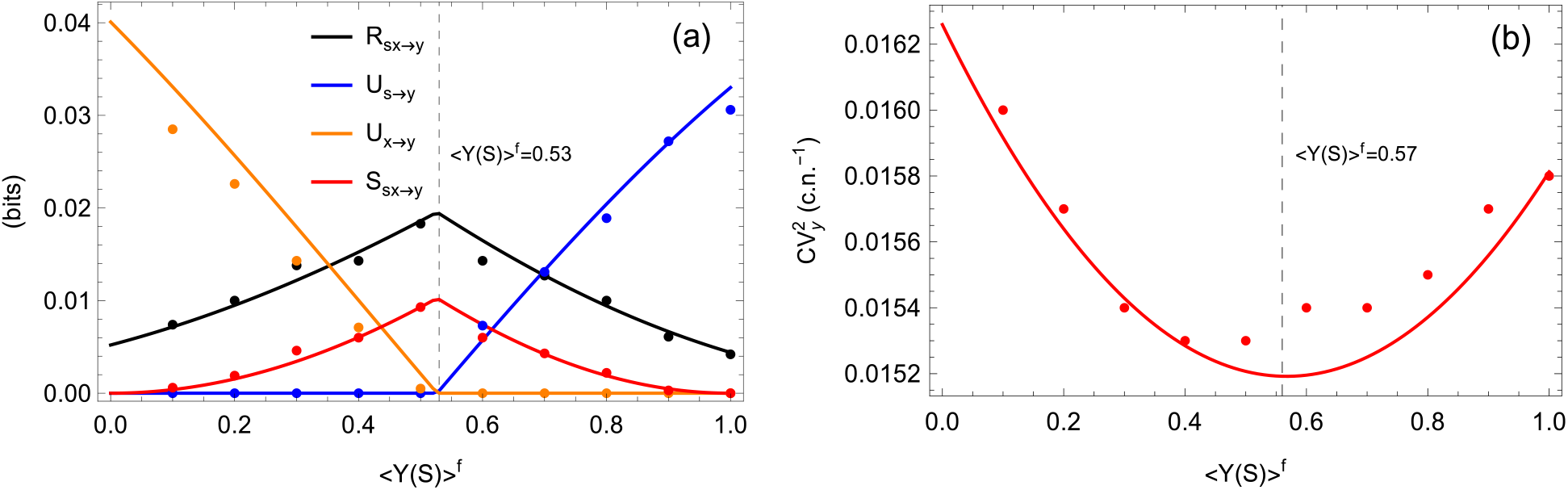
Panel (a) portrays the four PID elements representing different types of information transferred from the regulators to the output gene product, measured in the units of bits. Panel (b) shows the output gene expression noise measured using the squared coefficient of variation (CV) in the unit of c.n.^−1^. The solid lines are the approximate analytical results derived within the small-noise domain. The symbols are the steady-state average datasets from an ensemble of 5 × 10^4^ independent Gaussian time series generated from the exact Langevin stochastic simulation. The parameters common to both treatments are listed in Sec. II C. Besides, Δ*t* = 10^−1^ unit of time is used to generate the analytical and stochastic simulation-based results. The dashed gridlines demarcate the values of ⟨*Y* (*S*)⟩ ^*f*^ at which *R*_*sx*→*y*_ and *S*_*sx*→*y*_ are maximized in panel (a) and 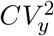 is minimized in panel (b). The mismatch between the analytical and the stochastic simulation-based datasets is due to the presence of intrinsic noise which is significantly generated from the discreteness of copy numbers of the gene products in the Langevin stochastic simulation.

Fig. 2(b) documents the response of the output gene expression noise in terms of 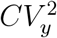. The concave-up profile has an opposite trend with respect to that of *R*_*sx*→*y*_ and *S*_*sx*→*y*_. As the master-regulator-influenced output production grows, the output noise tends to decrease until it becomes minimized for an approximately balanced C1-FFL (⟨*Y* (*S*)⟩ ^*f*^ = 0.57). Thereafter, the output noise begins to rise. For equal target production levels (⟨*Y* (*S*)⟩ and ⟨*Y* (*X*)⟩), the direct and indirect decoding processes of the input signal become equally strong. This establishes maximum control over the output variance leading to the minimization of the output noise. In the absence of such a balanced regulatory control, the output gene expression shows an increased noise level in either direction away from ⟨*Y* (*S*)⟩^*f*^ = 0.5. The exact positions of the redundant and synergistic information transfer maxima and the output noise minimum depend upon the intricate interplay among different systemic parameters. Consulting Figs. 2(a, b) together, we can appreciate the fact that the redundant and synergistic information transfers are appropriate predictors for the output gene expression noise. Looking at the values of ⟨*Y* (*S*)⟩ ^*f*^ at which *R*_*sx*→*y*_ and *S*_*sx*→*y*_ attain their peak values and output noise is minimized, it appears that the maximization of *R*_*sx*→*y*_ and *S*_*sx*→*y*_ can forecast subsequent minimization of the output noise and most importantly its forthcoming growth. Since, the output noise is an important quantifier of the corresponding gene expression stability which is invariably interlinked to the physiological fitness of a single cell, the redundant and synergistic information transfers may be considered as metrics of considerable biophysical utility. Due to the complex functional dependencies of the TE metrics and the output variance on the biochemical parameters, it is difficult to mathematically account for the relative positions of the redundant and synergistic information transfer maxima and the output noise minimum and the phenomenon demands further investigations. Nevertheless, we will test our current hypothesis in a reasonably broad and biochemically plausible parametric regime in the upcoming segments of our analysis. For ease of comparison, we will present the new datasets from the analytical computations due to the revised parameter choices alongside our control analytical data from Fig. 2.

### B. Effects of the temporal step Δ*t* on the dynamic information redundancy, synergy, and the output noise

The numerical value of the temporal step Δ*t* is of great significance in our analysis of the information dynamics. Eqs. (A.1a-A.1c) show that the synthesis and degradation reaction propensities at time point *t*, in general, are *f*_*z*_(*Z*_*t*_)Δ*t* and *µ*_*z*_*Z*_*t*_Δ*t*, respectively. On the other hand, the Gaussian noise processes, at time point *t* are 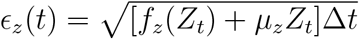. For the analytical purpose, we use the approximate steady-state average values (*f*_*z*_(⟨*Z*_*t*_⟩), *µ*_*z*_ ⟨*Z*_*t*_⟩) for the propensities and noise, a judicious choice for the computational tractability in sync with the small-noise approximation [24]. For the purpose of stochastic simulation, their exact time-dependent values are used. Changing the temporal step size affects the reaction probabilities and the random forcings linearly and sublinearly, respectively. Since, the analytical computations are performed within the small-noise regime, we choose a different value of Δ*t* which is smaller than the control value in Fig. 2 and by doing so ensure that the noise strengths in the dynamics decrease. It is to be reiterated that Δ*t* is also the temporal distance between the predictor and target variables. Hence, a decreased Δ*t* also decreases the numerical values of the predictor PID elements.

When we compare the PID elements analytically obtained using Δ*t* = 10^−1^ and 10^−2^ units of time in Fig. 3(a), we observe that their overall variational ranges in the former case are roughly 10 times the corresponding ranges in the latter case. Surprisingly, their variational trends with respect to ⟨*Y* (*S*)⟩ ^*f*^ is preserved in the Δ*t* transformation. When we examine the analytical profiles of the output noise in Fig. 3(b), it is found that 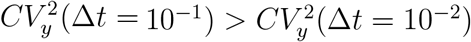, although in contrast with the PID metrics, the output noise does not undergo numerical change by any order of magnitude. We have compared the positions of the maxima and minima of the redundant and synergistic information transfers and the output noise, respectively, in Table I. From these results, we conclude that a reasonable alternative choice for Δ*t* does not alter the fact that the appearances of maxima of the redundant and synergistic information transfers precede the appearance of minimum of the output noise profile. Hence, the maximized redundant and synergistic information transfers can safely predict the minimization and the subsequent growth in the output noise level even when the temporal interval available for prediction (Δ*t*) is reasonably small.

**FIG. 3.**
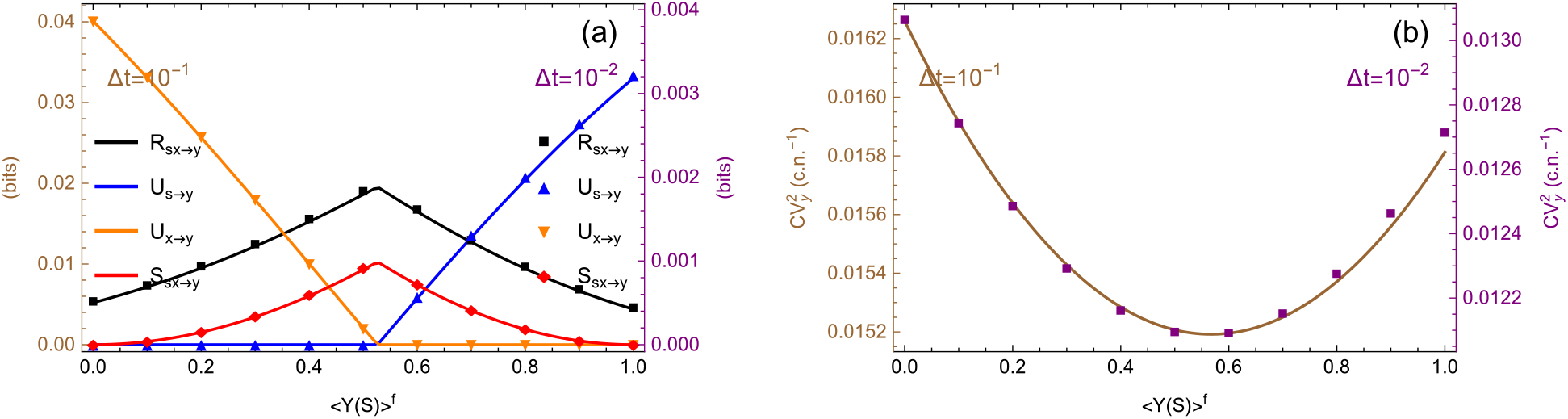
Panels [(a), (b)] compare {*R*_*sx*→*y*_, *S*_*sx*→*y*_} and 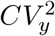, respectively, for two values of the temporal step Δ*t* = 10^−1^ (solid lines) and Δ*t* = 10^−2^ (symbols). All the datasets are obtained from the analytical calculations using the small-noise approximation. Differently colored vertical axes correspond to the reference scales for the statistical metrics obtained from different values of Δ*t*. The biochemical parameters used are identical to the case in Fig. 2.

**TABLE I.** The list compares the positions of the redundant and synergistic information transfer peaks and the output noise minima on the ⟨*Y* (*S*)⟩^*f*^ axis for different values of Δ*t* in Fig. 3.

| $\Delta t$ (unit of time) | $\{R_{sx \rightarrow y}, S_{sx \rightarrow y}\}_{\text{Max}}$ at $\langle Y(S) \rangle^f =$ | $CV_y^2\vert _{\text{Min}}$ at $\langle Y(S) \rangle^f =$ |
| --- | --- | --- |
| $10^{-1}$ | 0.53 | 0.57 |
| $10^{-2}$ | 0.52 | 0.56 |

### C. Effects of the input stability on the redundant and synergistic information transfers and the output noise

The set of degradation rate parameters {*µ*_*s*_, *µ*_*x*_, *µ*_*y*_} are the independently tunable parameters given the constraint of constant steady-state gene expression levels. These rate parameters are connected to the lifetimes of respective gene products through the relation *τ*_*z*_ = ln 2*/µ*_*z*_ and thus are the characteristic inverse timescales of the dynamics in Eqs. (1a-1c). As we increase the input relaxation rate parameter from the reference value *µ*_*s*_ = 0.1 to 1.0 and finally 5.0 min^−1^, Fig. 4(a) shows a decreasing *R*_*sx*→*y*_. The trend is opposite for *S*_*sx*→*y*_ in Fig. 4(b) and needs further in-depth mathematical explanation but for our present practical purpose, is irrelevant. It is also observed that with decreasing stability of the master-regulator, the redundant and synergistic information transfers are maximized at the cost of lesser amounts of the direct output production. In Fig. 4(c), the output noise, similar to the redundant information transfer, decreases with increasing *µ*_*s*_ but the positions of its minima do not follow any specific trend. Table II collates the positions of the maxima and minima on the ⟨*Y* (*S*)⟩^*f*^ axis for different values of *µ*_*s*_. The one important message conveyed by Table II is that the output noise minimization occurs after the redundant and synergistic information transfers are maximized in the system. Therefore, our hypothesis proposing the maximized synergistic and redundant information transfers as suitable predictors for the rise of the output gene expression noise is valid irrespective of the stability of the input signaling species that the C1-FFL motif may encounter in a fluctuating environment.

**FIG. 4.**
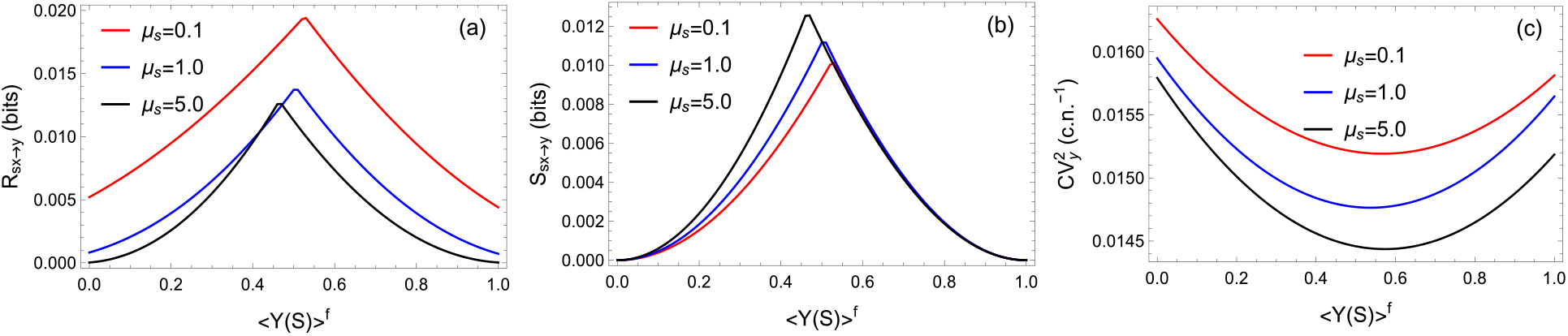
Each of the panels [(a), (b), (c)] shows variations in *R*_*sx*→*y*_, *S*_*sx*→*y*_, and 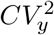, respectively, with respect to ⟨*Y* (*S*)⟩^*f*^ for *µ*_*s*_ = 0.1, 1.0, and 5.0 min^−1^. The datasets shown are obtained analytically using the small-noise approximation. All other independent parameters are kept fixed at their control values in Fig. 2.

**TABLE II.** Comparison table for the positions of the maximum and minimum points of the redundant and synergistic information transfers and the output noise, respectively, for different values of *µ*_*s*_ in Fig. 4.

| $\mu_s$ ($\text{min}^{-1}$) | $\{R_{sx \rightarrow y}, S_{sx \rightarrow y}\}\vert _{\text{Max}}$ at $\langle Y(S) \rangle^f =$ | $CV_y^2\vert _{\text{Min}}$ at $\langle Y(S) \rangle^f =$ |
| --- | --- | --- |
| 0.1 | 0.53 | 0.57 |
| 1.0 | 0.50 | 0.54 |
| 5.0 | 0.47 | 0.57 |

### D. Effects of the activation coefficients on the redundant and synergistic information transfer elements and the output gene expression noise

Next, we investigate the validity of our core hypothesis when the activation coefficients: *K*_*sx*_, *K*_*sy*_, and *K*_*xy*_ are set at an operating point other than the half-maximal point as in the control data in Fig. 2. The TF-promoter binding strength is measured by the inverse of the activation coefficient and hence an increased *K*_*···*_ signifies that the transcriptional link is weakened. Depending upon the value of the activation coefficient with respect to the associated TF copy number, the (non)linear nature of the input Hill function may change. To exemplify, with *K*_*sx*_ *≫ S*, the synthesis rate of X becomes 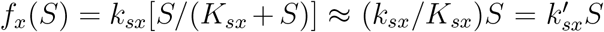. Hence, the transcriptional regulation becomes linear in *S* (first-order kinetics) with a reduced maximum production rate 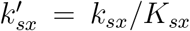. On the other hand, with *K*_*sx*_ *≪ S*, an approximate zero-order kinetics follow as *f*_*x*_(*S*) ≈ *k*_*sx*_. The half-maximal activation point due to *K*_*sx*_ = *S* retains a sufficient amount of (non)linearity which suits the requirements of PID. Notably, the MMI PID is valid when the dynamics under investigation is either linear or linearized. Taking this into our consideration, we intend to vary various *K*_*···*_ applying reasonably small changes around their half-maximal values. The activation coefficient can be changed by means of altering the gene promoter sequences so that the respective promoter’s chemical affinity towards its input TF is changed [2]. For *K*_*···*_ = ⟨*Z*⟩ ± 20% ⟨*Z*⟩ = 1.2 ⟨*Z*⟩, 0.8 ⟨*Z*⟩, the various approximate steady-state values of the Hill functions associated with the independent reaction channels become ≈ 0.455, 0.556, respectively.

When we change *K*_*sx*_ taking *K*_*sx*_ = ⟨*S*⟩ = 100 as our datum line and keeping *K*_*sy*_ = ⟨*S*⟩ and *K*_*xy*_ = ⟨*X*⟩, Fig. 5(a) shows that an increased (decreased) *K*_*sx*_ lifts (drops) the redundant information transfer. Similar trends are also visible in the output noise level portrayed in Fig. 5(e) whereas the synergistic information transfer is robust to the altered *K*_*sx*_ as shown in Fig. 5(c). When the direct and indirect transcriptional links regulating the output gene expression are subjected to similar types of change while keeping *K*_*sx*_ = ⟨*S*⟩, the data projects that increasing (decreasing) {*K*_*sy*_, *K*_*xy*_} results in increased (decreased) levels of 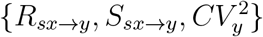 as profiled in Figs. 5(b, d, f), respectively. These results may be explained by taking into consideration the fact that with reduced activation coefficients, the binding affinities become higher or in other words, the input TFs bind strongly with their target promoters and such tight binding events lower the fluctuations in the transcriptional activation reactions. Therefore, the levels of the output noise and most of the redundant and synergistic information transfers drop in magnitudes. The changes in relevant statistical metrics around the reference half-maximal data (red solid lines) in Figs. 5(b, d, f) are larger in magnitudes in comparison with their counterparts in the variable *K*_*sx*_ case, i.e., Figs. 5(a, c, e), respectively. These datasets show that the transferred informations and the output noise are less susceptible to the change in the binding affinity of S → X than in S → Y and X → Y. This is possibly due to the fact that the former link is static in terms of fixed production level ⟨*X*⟩ whereas the latter two edges are sufficiently dynamic because of varying ⟨ *Y* (*S*)⟩ and ⟨*Y* (*X*)⟩ levels. The data in Fig. 5(c) showing an almost identical response of *S*_*sx*→*y*_ towards different values of *K*_*sx*_ need further interpretation in terms of detailed changes in the analytical expressions of the single- and two-time-point (co)variances and also possibly in the TE metrics. But due to the presence of the nonlinear logarithm function, such fine-grained mathematical explanation is difficult to achieve. Fortunately, our main concerns in these measurements involve the antagonistic trends of {*R*_*sx*→*y*_, *S*_*sx*→*y*_} with respect to 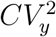 and the appearances of the information maxima followed by the noise minima. The values of ⟨*Y* (*S*)⟩^*f*^ for which the redundant and synergistic information transfers are maximized and the output noise is minimized for different numerical values of the activation coefficients, are presented in Table III. These datasets categorically suggest that even when the operating points on various activation curves are shifted from the respective half-maximal points, *R*_*sx*→*y*_ and *S*_*sx*→*y*_ attain their peaks before 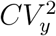 reaches its lowest value. Hence, these two PID flavors again become enabled to forecast the growth in the output gene expression fluctuations.

**FIG. 5.**
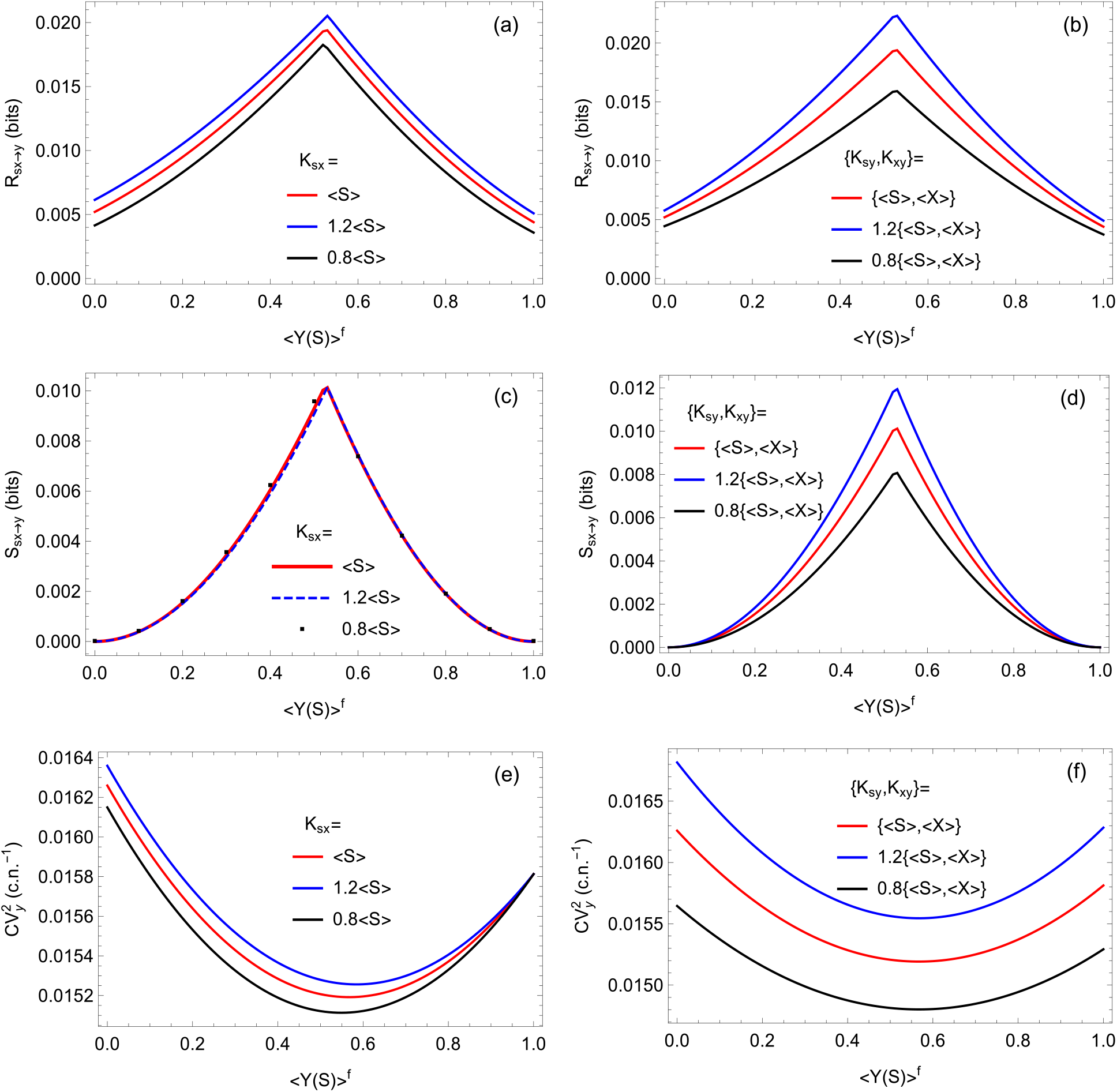
Panels [(a), (c), (e)] show the profiles of *R*_*sx*→*y*_, *S*_*sx*→*y*_, and 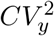, respectively, for different values of *K*_*sx*_. Panels [(b), (d), (f)] contain the plots for these three metrics in the same order but for different values of *K*_*sy*_ and *K*_*xy*_. To showcase a reasonable number of interesting parametric realizations, the constraint of *K*_*sy*_ = *K*_*xy*_ is maintained. Each of the three types of activation coefficients is assigned values such that two points around the respective half-maximal points on different activation curves may be accessed. Apart from the half-maximal case, the other two values for each of the activation coefficients are assigned according to the prescription: *K*_*···*_ = ⟨*Z*⟩ ± 20% ⟨*Z*⟩. Other independent parameters and population levels are kept unchanged from Fig. 2. All the datasets are analytical results obtained within the small-noise domain.

**TABLE III.** The values of ⟨*Y* (*S*)⟩^*f*^ at which the redundant and synergistic information transfers are maximized and the output noise is minimized for different values of the activation coefficients are listed. Only those activation coefficients, whose variations are investigated, have been explicitly mentioned and the table should be consulted along with Fig. 5. Other activation coefficients are kept fixed at their respective half-maximal values. The activation coefficients are expressed in terms of the population levels: ⟨*S*⟩ = ⟨*X*⟩ = ⟨*Y*⟩ = 100.

| Activation coefficients | $\{R_{sx \rightarrow y}, S_{sx \rightarrow y}\}\vert _{\text{Max}}$ at $\langle Y(S) \rangle^f =$ | $CV_y^2\vert _{\text{Min}}$ at $\langle Y(S) \rangle^f =$ |
| --- | --- | --- |
| $K_{sx} = \langle S \rangle$ | 0.53 | 0.57 |
| $K_{sx} = 1.2\langle S \rangle$ | 0.53 | 0.58 |
| $K_{sx} = 0.8\langle S \rangle$ | 0.52 | 0.55 |
| $\{K_{sy}, K_{xy}\} = \{\langle S \rangle, \langle X \rangle\}$ | 0.53 | 0.57 |
| $\{K_{sy}, K_{xy}\} = 1.2\{\langle S \rangle, \langle X \rangle\}$ | 0.53 | 0.57 |
| $\{K_{sy}, K_{xy}\} = 0.8\{\langle S \rangle, \langle X \rangle\}$ | 0.53 | 0.57 |

### E. Effects of differential gene expressions of the regulators and the output species on the redundant and synergistic information transfers and the output noise

The gene expression levels of the predictor and target species are important factors determining how much redundant and synergistic informations are transferred from the input to the output level in the motif and the extent to which the target’s fluctuations are altered? We bring in significant asymmetry between the populations of the information sources and target to demonstrate this point while keeping the parametric realization in Fig. 2 as our reference. Since, both S and X are the information providers, we retain their populations identical to each other (⟨*S*⟩ = ⟨*X*⟩ = 100) while that of their target is either increased or decreased significantly.

Fig. 6(a) suggests that when the regulators predict the target species with a population pool twice as strong as the sources (⟨*Y*⟩ = 200), both *R*_*sx*→*y*_ and *S*_*sx*→*y*_ increase with respect to the reference datasets of Fig. 2(a). On the output noise side in Fig. 6(c), due to its population-normalized definition 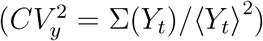, increased ⟨*Y*⟩ (or equivalently ⟨*Y*_*t*_⟩) invariably results in decreased output fluctuations. In the opposite scenario, i.e., regulators predicting their target with a population half of that of the former species (⟨*Y*⟩ = 50), the redundant and synergistic information transfers and the output noise level are decreased and increased, respectively, with regards to the designated control datasets [see Figs. 6(b, d)]. The opposite trend of the output noise profile with respect to the redundant and synergistic information transfers reappears in the present parameterization and moreover ensures that the relevant information transfer elements maximize before the output fluctuations are minimized. The specific positions of different maxima and minima are listed in Table IV. Therefore, even with the asymmetrical production machineries of the regulators and their enslaved gene product, the maximized redundant and synergistic information transfers can efficiently predict the growth of the output noise.

**FIG. 6.**
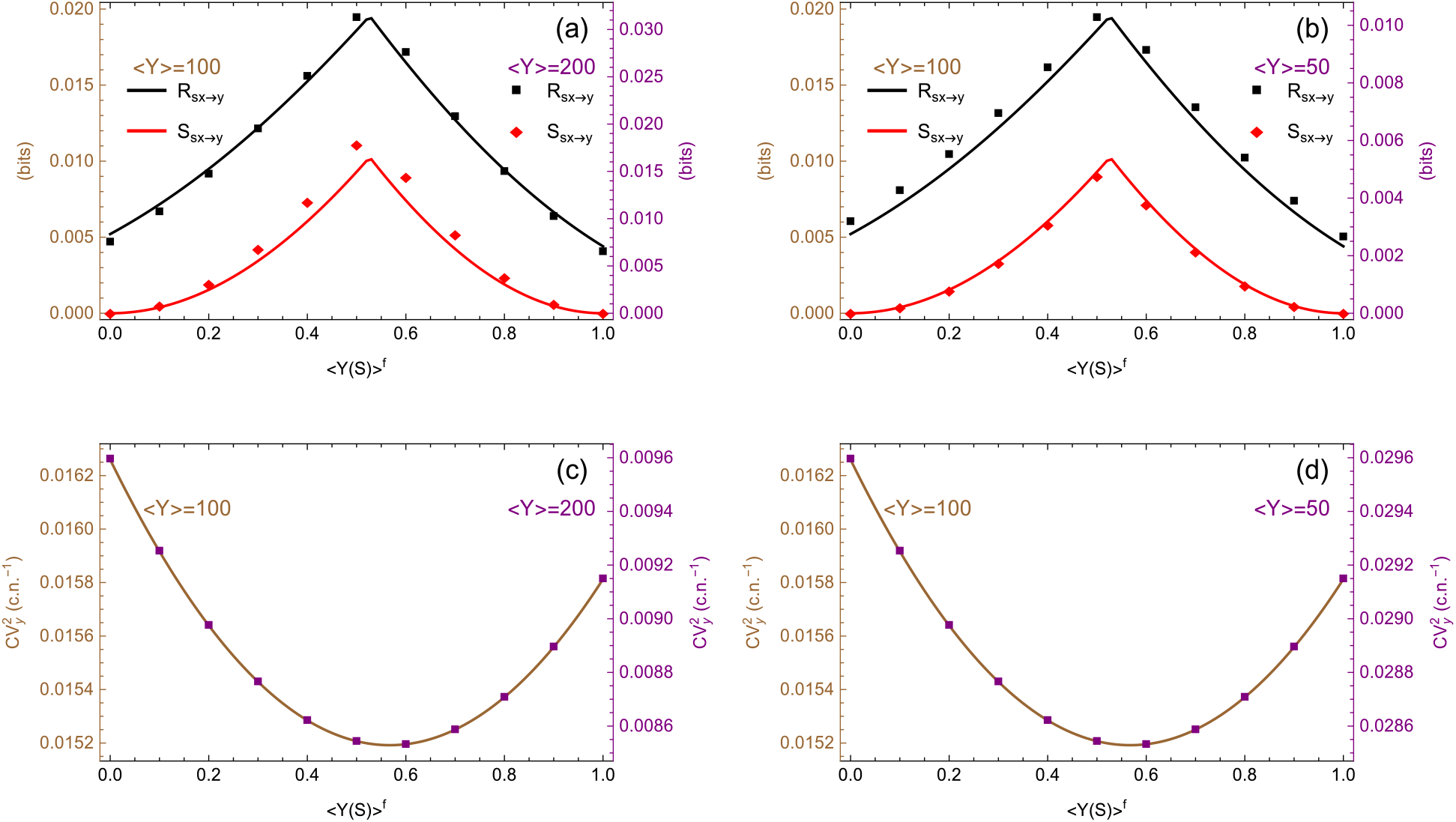
Panels [(a), (c)] compare {*R*_*sx*→*y*_, *S*_*sx*→*y*_} and 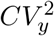, respectively, for ⟨*Y*⟩ = 200 (symbols) with the reference data generated for ⟨*Y*⟩ = 100 (solid lines). Panels [(b), (d)] do similar comparisons for the data due to ⟨*Y*⟩ = 50 (symbols). The regulator abundances are kept identical to the previous cases, i.e., ⟨*S*⟩ = ⟨*X*⟩ = 100. The constraint of ⟨*Y* (*X*)⟩ = ⟨*Y*⟩ − ⟨*Y* (*S*)⟩ is still in place so are other independent choices of different parameters. The datasets are derived from analytical computations under the aegis of small-noise approximation.

**TABLE IV.** The values of ⟨*Y* (*S*)⟩^*f*^ corresponding to the maxima of the redundant and synergistic information transfers and the minima of the output noise for different output gene expression levels are tabulated. The regulator population levels are kept at ⟨*S*⟩ = ⟨*X*⟩ = 100. The profiles presented in Fig. 6 should be consulted along with these datasets for various extrema.

| Output gene expression level ($\langle Y \rangle$) | $\{R_{sx \rightarrow y}, S_{sx \rightarrow y}\}\vert _{\text{Max}}$ at $\langle Y(S) \rangle^f =$ | $CV_y^2\vert _{\text{Min}}$ at $\langle Y(S) \rangle^f =$ |
| --- | --- | --- |
| 100 | 0.53 | 0.57 |
| 200 | 0.525 | 0.565 |
| 50 | 0.52 | 0.56 |

## IV. CONCLUSION

In summary, we have studied the information transfer and output noise management capacities of various manifestations of a generic C1-FFL motif. For fixed steady-state abundances of the three constituent gene products, we have invoked the small-noise approximation within a Gaussian framework and applied the principle of MMI PID to the time-discrete MVAR dynamics. This has helped us to quantify the redundant, synergistic, and unique flavors of information transfer from the upstream species to the final gene product. For an approximately balanced C1-FFL (equally strong direct and indirect decoding strengths), these two types of dynamic information dominate their unique counterparts which become vanishingly small as the redundant and synergistic information transfers are maximized in the system. Hence, with appreciably strong direct and indirect regulations of the target (output) species, the regulators cease to independently predict their target and only act together.

We have demonstrated that the redundant and synergistic information transfers are antagonistically related to the output noise level. Moreover, these information transfer elements are maximized before the minimization and subsequent growth of the output noise. Depending upon these findings, we hypothesize that the redundant and synergistic information transfer elements may be referred to as potential predictors (indicators) for the rise in the fluctuation level associated with the output gene expression. Since, the output gene product may serve as an essential enzyme/protein/TF for various crucial physiological purposes of a single cell, a predictive control over its fluctuation is of utmost importance for the cell’s survival in a changing environment. We have supported our main piece of the analytical result with data generated from the exact procedure of Langevin stochastic simulation coupled with sufficient ensemble averaging over a large number of independently generated Gaussian time series.

The order in which the redundant and synergistic information transfer maxima and the output noise minimum appear with respect to changing ⟨*Y* (*S*)⟩^*f*^ remains a puzzle both at the mathematical and biochemical levels of understanding. We hope that further research will be able to demystify the phenomenon in its entirety. Nevertheless, we have tested our core hypothesis under sufficiently varied choices for different parameters. We have found out that our main result stands firmly when: (i) the time duration available to the regulators for predicting their target is reduced by an order-of-magnitude, (ii) the input species’ stability is significantly modified, (iii) various activation coefficients are tuned in and around their individual half-maximal operating points, and (iv) the population level of the predicted output is made to be double/half of the individual populations of its two upstream predictor species. Thus, our main finding regarding the predictive capacities of the redundant and synergistic information transfers targeting the output gene expression noise is pretty robust with respect to reasonable variations of the relevant biochemical parameters conforming to the small-noise approximation.

In the absence of a standardized MMI PID technique for more than three stochastic variables, the generic C1-FFL network with tunable transcriptional control over the output gene product is an ideal model system to systematically investigate the response of different flavors of information transfer to various important biochemical parameters. Previously, the principle of MMI PID has been applied to the two-dimensional Ising model and the findings have categorically demonstrated that the redundant and synergistic information transfer terms may act as warning signals for the transition from a disordered to an ordered state [29]. Our present hypothesis should be further tested for its generality, e.g., when the biochemical system involves transitions between different steady states. Since, the prescription of MMI PID is valid for either linear or reasonably linearized systems, it remains an open question whether the present theoretical approach can be safely applied to such nonlinear biochemical transitions. The situation is further complicated due to the presence of large fluctuations in and around the point of transition and therefore the present analytical treatment using the small-noise approximation will be defunct in tackling such problems. Explorations along these lines of research are supposed to generate new insights about the real-world applications of the multivariate information theory.

## ACKNOWLEDGMENTS

Md SAM is supported by DST, Govt. of India, through INSPIRE research fellowship (DST/INSPIRE Fellowship/2018/IF180056). Md SAM and AB acknowledge Bose Institute, Kolkata for research support.

## Appendix Computing the time-discrete variances and covariances

### Analytical expressions using the small-noise approximation

The Langevin equations (1a-1c) are in the continuous-time domain. In this section, we use the MVAR process to calculate TEs between different nodes of the reference C1-FFL network [31– 33]. To do so, we discretize Eqs. (1a-1c) considering the gene expression dynamics with a finite temporal step/resolution (Δ*t*). In this time interval, changes in the population level of species S is *S*_*t*−1_ → *S*_*t*_ where *t* − 1 symbolically denotes the immediate past (i.e., Δ*t* time behind) time point of *t*. The consequent rate of change in copy number state is 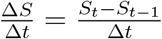. The equivalence between the continuous and discrete versions is through the relation: 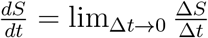. Hence, the time-discrete Langevin equations are approximations to their continuous counterparts. The resulting discretized dynamical system of equations is as follows:

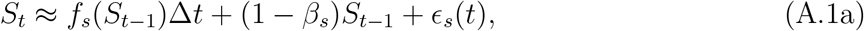

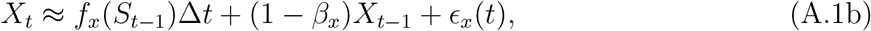

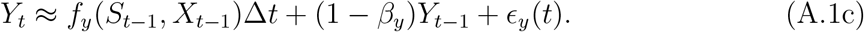

Here, we have simplified our anaytical treatment by assuming *ϵ* _*z*_(*t* − 1) ≈ *ϵ* _*z*_(*t*). Since, our forthcoming analysis is intended to be at steady state, approximating noise terms from neighbouring time points to be equal to each other is a useful simplification for the analytical purpose. This is based upon the assumption that the steady-state copy numbers do not change significantly between two neighbouring time points, i.e., ⟨*Z*_*t*−1_⟩ ≈ ⟨*Z*_*t*_⟩. Since, the noise strenghts are functions of the copy numbers, we can safely use approximately equal noise strengths for time points *t* and *t*−1. However, while executing the exact stochastic simulation, such an approximation is not tenable because the system’s simulated dynamics also involve transient states. Additionally, within a finite time-span Δ*t*, depending upon the number of reactions happening, *Z*_*t*−1_ ≈ *Z*_*t*_ may not hold for all the time points (even at steady state) each time an independent Gaussian stochastic time series is generated in repeated time-course simulation experiments. For fixed Δ*t, β*_*z*_ = *µ*_*z*_Δ*t* represent the inverse timescales in dimensionless forms for our time-discrete dynamical system. Therefore, the rescaled random forcing terms for this discretized system are 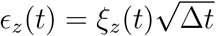. Using the properties of *ξ*_*z*_(*t*) from Sec. II A, we note that ⟨*ϵ*_*z*_(*t*)⟩ = 0 and ⟨*ϵ*_*z*_(*t*)*ϵ*_*z* ′_ (*t′*) = ⟨|*ξ*_*z*_(*t*)|^2^⟩Δ*tδ*_*zz*′_*δ*(*t* − *t′*). The steady-state ensemble-averaged noise strengths in Eqs. (A.1a-A.1c) are related to their counterparts in Eqs. (1a-1c) through the relation ⟨| *ϵ*_*z*_(*t*)|^2^⟩= ⟨|*ξ*_*z*_(*t*)|^2^⟩Δ*t*. Ensemble-averaging Eqs. (A.1a-A.1c) at steady state, we obtain:

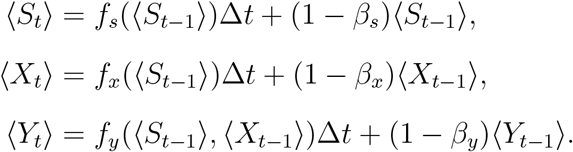

Where, we have utilized the fact that ⟨ *ϵ*_*z*_(*t*)⟩ = 0. Now, at time point *t*, the copy number fluctuations around the respective steady-state mean values are:

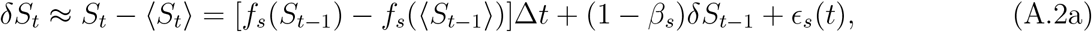

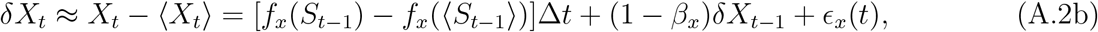

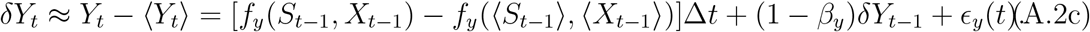

Here, we have invoked the small-noise approximation [40, 41, 51, 52], according to which, at steady state, the copy number fluctuations of order higher than the linear one is appreciably smaller and can be neglected when compared to the steady-state mean copy numbers. To paraphrase mathematically, this approximation implies that *δZ*_*t*_/ ⟨*Z*_*t*_⟩ ≪1 where *Z*_*t*_ ∈ {*S*_*t*_, *X*_*t*_, *Y*_*t*_}. The condition is similarly valid for the immediate past time point. Hence, we can rewrite the steady-state perturbations in the input functions due to the infinitesimal changes, i.e., {*δS*_*t*−1_, *δX*_*t*−1_} → 0 in the following manner:

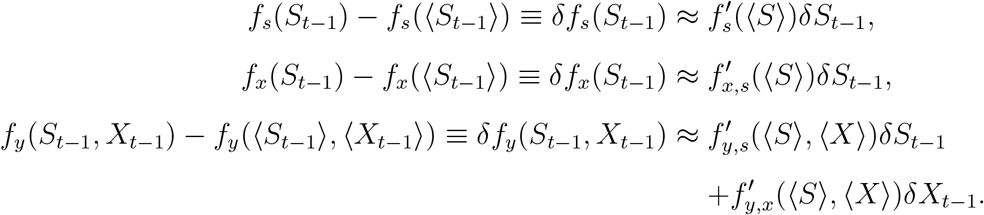

Here, we have taken recourse to: ⟨*S*_*t*−1_⟩≈ ⟨*S*_*t*_⟩≡ ⟨*S*⟩ and ⟨*X*_*t*−1_⟩≈ ⟨*X*_*t*_⟩≡ ⟨*X*⟩. Here, 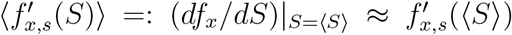 and so on for other derivatives of the input functions. In our analytical calculations, performed within the small-noise domain, we have used a couple of judicious steady-state approximations, e.g., ⟨*f*_*x*_(*S*)⟩≈ *f*_*x*_(⟨*S*⟩) and ⟨*f*_*y*_(*S, X*)⟩≈ *f*_*y*_(⟨*S*⟩, ⟨*X*⟩). Extending this approximation to the derivatives, we have: 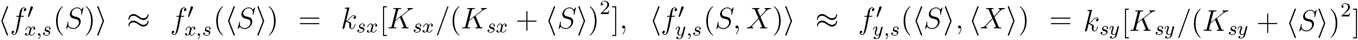, and 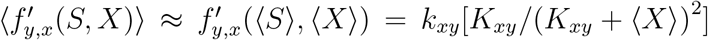. Nevertheless, while executing the stochastic Langevin simulation, we have used actual copy numbers instead of their steady-state values in the input functions/reaction propensities [24]. Similar approximations at the previous time point lead us to ⟨*f*_*s*_(*S*_*t*−1_)⟩≈ *f*_*s*_(⟨*S*_*t*−1_⟩), ⟨*f*_*x*_(*S*_*t*−1_)⟩ ≈ *f*_*x*_(⟨*S*_*t*−1_⟩), ⟨*f*_*y*_(*S*_*t*−1_, *X*_*t*−1_)⟩ ≈ *f*_*y*_(⟨*S*_*t*−1_⟩, ⟨*X*_*t*−1_⟩). For the purpose of stochastic simulation, we have used dynamic input functions, i.e., *f*_*z*_(*Z*_*t*−1_) instead of their steady-state values: ⟨*f*_*z*_(*Z*_*t*−1_)⟩or, as in our analytics, *f*_*z*_(⟨*Z*_*t*−1_⟩). For the details of the stochastic simulation method, see the last part of this section. Rewriting the set of Eqs. (A.2a-A.2c), we get:

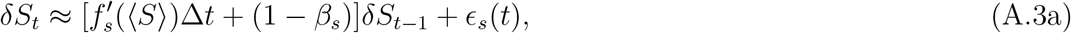

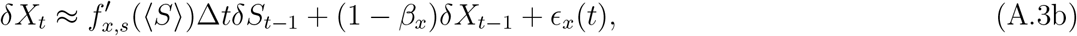

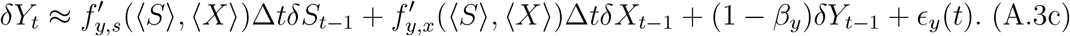

Eqs. (A.3a-A.3c) in matrix form stand as:

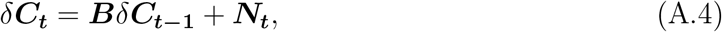

where,

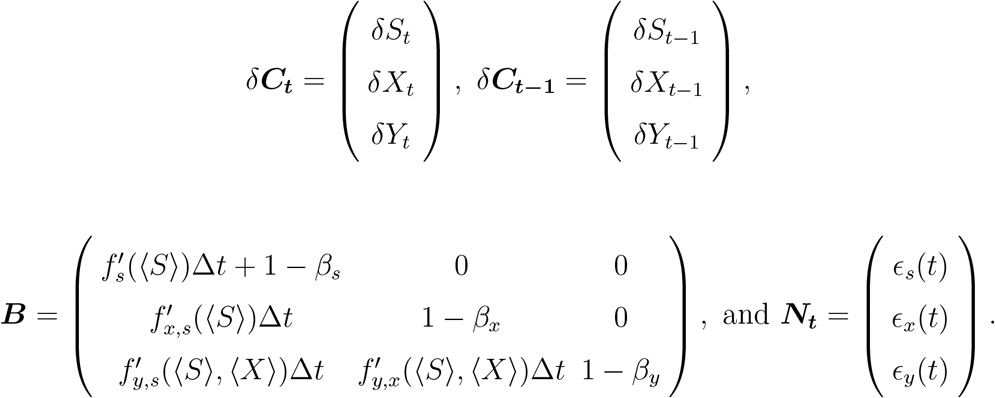

By multiplying Eq. A.4 by its transposed counterpart and then performing the steady-state ensemble-averaging, we arrive at:

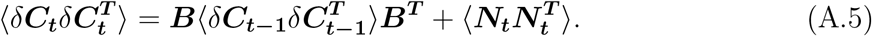

Since, the mean squared deviations of the copy numbers are approximately stationary at steady state, they are approximately equal to each other at successive steady-state time points. Hence, we can write:

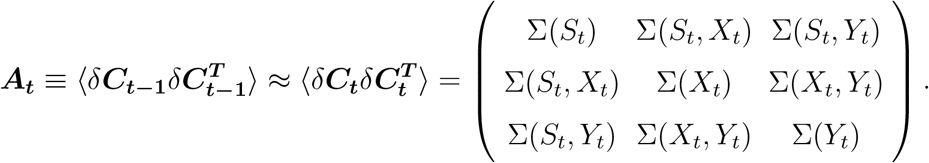

We denote the matrix incorporating the steady-state ensemble-averaged noise strengths as 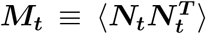. In the absence of cross-species noise-correlations, ***M***_***t***_ is a diagonal matrix with elements ⟨| *ϵ*_*z*_(*t*)|^2^⟩. Finally, we have a steady-state fluctuation-dissipation type of equation:

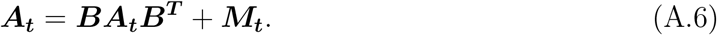

Solving Eq. A.6, we arrive at the closed-form analytic expressions of the steady-state copy number variances and covariances. Here for notational simplicity, we have denoted the steady-state derivatives of different input functions as: 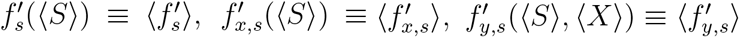, and 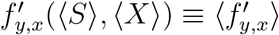.

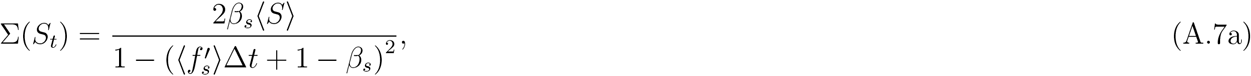

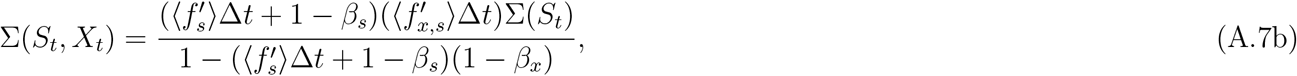

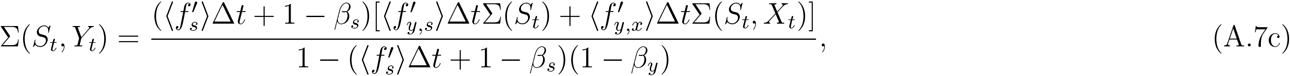

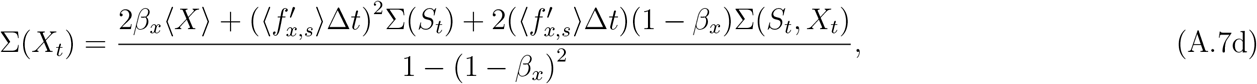

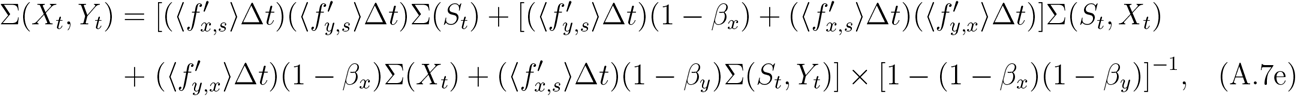

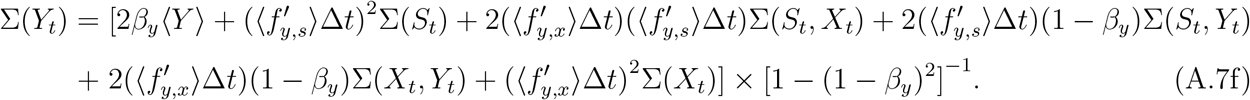

In the next step, we compute the one-time-lagged covariances by replacing *t* by *t* + 1 in Eq. A.4. We have again used the simplifying assumption ***N***_***t*+1**_ ≈ ***N***_***t***_ considering our steady-state analytical requirements. This leads us to the following:

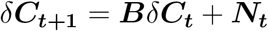

By multiplying the above equation with 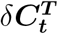 and upon subsequent steady-state ensemble-averaging, we get:

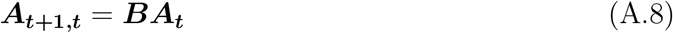

Here, we have used the fact that 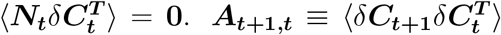 is the one-time-lagged covariance matrix, individual elements of which, can be obtained by solving Eq. A.8. The closed-form analytic expressions are listed below:

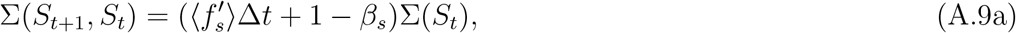

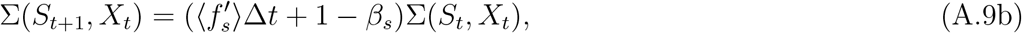

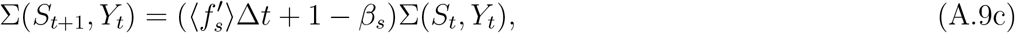

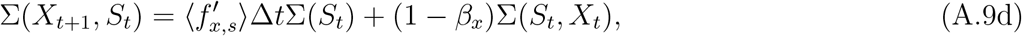

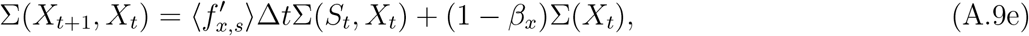

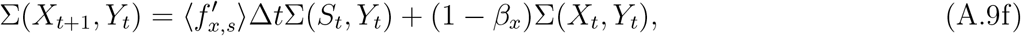

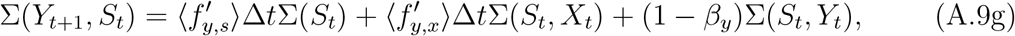

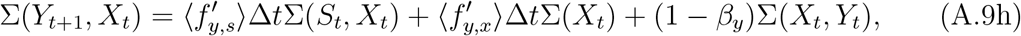

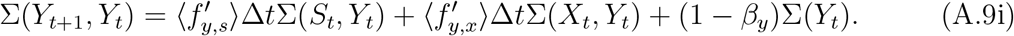

Here, 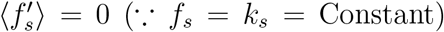. According to the prescription in Ref. [32], the partial variances can be written as:

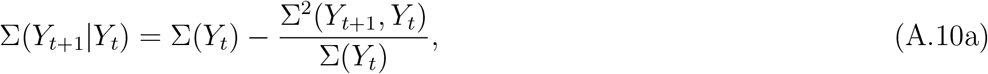

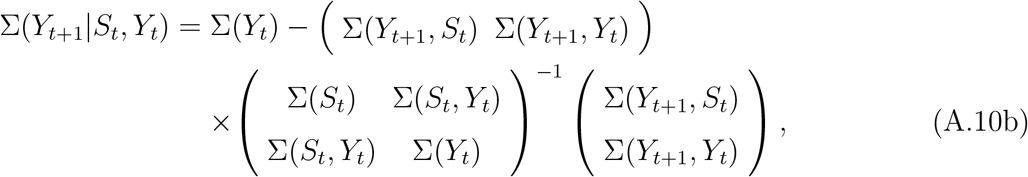

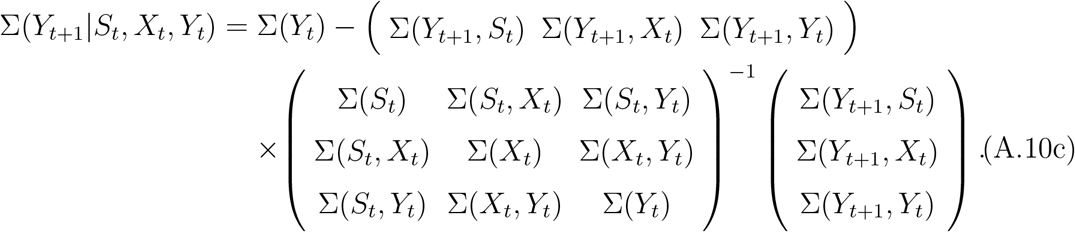

The working formula for Σ (Y_*t*+1_|X_*t*_; Y_*t*_) can be deduced similarly.

### Computations using the exact Langevin stochastic simulation method

Expressions (A.7a-A.7f) and (A.9a-A.9i) are essential ingredients to analytically evaluate the PID elements and output noise within the small-noise regime. To validate the approximate results, we further execute stochastic simulation using Eqs. (A.1a-A.1c). Unlike the analytical case, we use exact copy numbers (*Z*_*t*−1_) instead of their steady-state average values (⟨*Z*_*t*−1_⟩) in the input functions *f*_*z*_(· · ·) as well as the degradation terms in the right-handside of these equations. Similarly, the noise strengths are utilized as functions of actual copy numbers and not their steady-state average values. As mentioned earlier, for the purpose of stochastic simulation, we maintain *ϵ*_*z*_(*t* − 1) ≠ *ϵ*_*z*_(*t*). Therefore, for the stochastic trajectories, we have |*ϵ*_*z*_(*t* − 1)|^2^ = |*ξ*_*z*_(*t* − 1)|^2^Δ*t* = [*f*_*z*_(*Z*_*t*−1_) + *µ*_*z*_*Z*_*t*−1_]Δ*t* as the noise strength operating at time point *t* − 1. For each of the three species, the associated Gaussian noise terms with zero means and unit variances are generated from the random number generating function *randn* in MATLAB. Hence, the modified random forcing in the simulated version of Eqs. (A.1a-A.1c) appears as 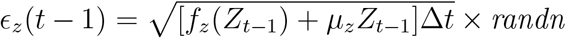. The choice for the numerical value of Δ*t* is crucial in the stochastic simulation for a couple of reasons. As mentioned before, the modified noise strength at time point *t*, i.e., |*ϵ*_*z*_(*t* − 1)|^2^ ∝ Δ*t*. Hence, a larger Δ*t* will evidently increase the stochasticity in the system and the simulated values of our metrics of interest will cease to be in sync with their analytical counterparts which are evaluated within the small-noise regime. On the other hand, the reaction propensities for different synthesis and degradation events happening in between time points *t* − 1 and *t* are *f*_*z*_(*Z*_*t*−1_)Δ*t* and *µ*_*z*_*Z*_*t*−1_Δ*t*, respectively. Thus, a smaller Δ*t* will make these reactions less probable in each time step and the predictability of future target states by the present source states will be hampered. As a result, the PID measures will be significantly reduced in their magnitudes. In fact, it can be analytically shown that with Δ*t* → 0, the variances and covariances obtained in Eqs. (A.7a-A.7f) reduce to their corresponding continuous-time versions derived in Ref. [35]. Therefore, one has to choose a reasonable value for Δ*t* keeping in mind both of these trade-offs/constraints. For the present case, we simulate the exact versions of the discretized Langevin Eqs. (A.1a-A.1c) for sufficiently long time *t*_*f*_ (200 units of time) so that (i) the system reaches a steady state and (ii) using an appropriate Δ*t* (10^−1^ unit of time), we can ensure that the system evolves via a sufficiently large number of iterations (*t*_*f*_ */*Δ*t* = 2000). Sec. III B analytically demonstrate that even with a smaller value of Δ*t* and consequently reduced values of the PID elements, our core hypothesis remains intact.

In this way, for each of the values of *k*_*sy*_ (*k*_*xy*_), we obtain a large number of independent Gaussian time series (5 ×10^4^) for the dynamical variables. The rationale/constraint through which the direct and indirect synthesis rate parameters of the output species are tuned is specified in Sec. II C. In the next stage, the necessary steady-state first and second moments (both single-time-point and one-time-lagged) of the copy number distributions are obtained considering the time points *t*_*f*_ and *t*_*f*_ −1 by averaging over the previously mentioned ensemble of time series. These are subsequently used to compute the single- and two-time-point (co)variances for the framework of PID. The averaged imprint of noise at steady state *t* = {*t*_*f*_, *t*_*f*_ −1} from this large collection of independent time series corresponds to the mean-field usage of ⟨| *ϵ*_*z*_(*t*)|^2^⟩in our approximated analytical calculations [see ***M***_***t***_ in Eq. A.6]. For an individual stochastic variable, one can conceptualize these independent stochastic time series as the gene expression profiles from individual single cells in a clonal population. Otherwise, these time traces may be interpreted as gene expression profiles gathered from a single cell in repeated time-course experiments with identical initial conditions and other constraints. Due to the probabilistic nature of biochemical reactions in a single cell, these gene expression profiles are all different from each other, a phenomenon which is popularly known as *isogenic heterogeneity* [9]. The Langevin stochastic simulation, in essence, mimics this biological reality. Since, this process involves none of the simplifying steps applied in our previous analytical treatment, the obtained statistical metrics are exact here. Sec. III A presents our main results using both analytical and stochastically simulated data for the PID elements and the output noise so that a meaningful and predictive comparison may be done.

